# Exploring phenotype-related single-cells through attention-enhanced representation learning

**DOI:** 10.1101/2024.10.31.619327

**Authors:** Qinhua Wu, Junxiang Ding, Ruikun He, Lijian Hui, Junwei Liu, Yixue Li

**Author notes:** Correspondence;, Preferred corresponding institute: Guangzhou National Laboratory.

## Abstract

The scope of atlas-level single-cell investigations reveals the pathogenesis and progression of various diseases. Accurate interpretation of phenotype-related single-cell data necessitates the pre-definition of single-cell subtypes and the identification of their abundance variations for downstream analysis. In this context, biases from batch correlation and the selection of clustering resolutions can significantly impact single-cell data analysis and result interpretation. To strengthen the associations across single cells in each sample and their clinical phenotype, and to enhance single-cell exploration by integrating cell and gene-level information. This study proposes a method to learn phenotype-related sample representations from single cells via the attention-based multiple instance learning (AMIL) mechanism. This approach incorporates gene expression profiles from each single cell for sample-level clinical phenotype prediction. By integrating deep learning interpretation methods and phenotype-specific single-cell attention weights across sample groups, this method highlights critical gene programs and cell subtypes that mostly contribute to the sample-level clinical phenotype, and facilitate mechanistic exploration. Using single-cell atlases from COVID-19 infected patients and age-related healthy human blood, we demonstrate that this method can accurately predict disease severity and age-related phenotypes. Additionally, variations in cellular attention reflect the underlying biological mechanisms associated with these phenotypes. This method proposes a supervised framework for single-cell data interpretation and can be further adapted for other atlas-level clinical phenotype analyses.

## Introduction

The development of single-cell RNA-seq methods allows for the profiling of gene expression at the level of individual cells, which provides valuable insights into the diverse states of single cells across tissue differentiation during development, aging, disease progression, and tumor drug resistance^1^. These single-cell atlases aim to explore the underlying links between disease phenotypes and single-cell targets. Despite its potential, the analysis of scRNA-seq datasets confronts significant challenges, mostly related to the high sparsity and noise in these data. Additionally, current computational methods often rely on single-cell data integration, clustering, and differential distribution analyses to identify disease-related single-cell subsets, and then to understand the distinct cell response mechanisms and gene regulation within these cells for understanding disease phenotypes^2^. However, these analytical processes are frequently hampered by data discrepancies across different platforms, batch effects from various experiments, and the performance limitations of single-cell integration algorithms. These issues can significantly distort the clustering and annotation of single-cell subsets, ultimately affecting downstream functional analysis^3^. In addition, unsupervised clustering of single-cell data tends to overlook common regulatory mechanisms among cell subsets, making it challenging to identify overarching disease-related features. Therefore, there is a critical need to accurately identify disease phenotype-specific cells and their gene regulatory relationships within single-cell datasets.

To strengthen the integration of single-cell data with clinical phenotypes, various methods have been developed to predict clinical outcomes based on single-cell gene expression data. The CloudPred method uses mixture of Gaussians to model single-cell data and extracts patient-specific features for phenotype prediction. The variations in the means and covariances of the predicted model offer valuable insights for biological interpretation^4^. scPheno learns the joint distribution of cell and disease phenotypes from scRNA-seq data to predict clinical outcomes^5^. ScRAT employs sample mixup and self-attention modules to generate fixed-sized embeddings of pseudo-samples for phenotype classification^6^. These approaches leverage deep learning and statistical methods to address the challenges posed by the sparsity and noise in single-cell gene expression data, as well as the variation in cell size across different samples. Additionally, other methods have been designed to directly explore associations within phenotypes and single cells. DA-seq quantifies the prevalence of distinct states within cellular neighborhoods, identifying novel differentially abundant cell clusters for biological interpretation^7^. Similarly, Dann et al. conducted atlas-level analyses to characterize disease-specific cellular distributions in comparison to healthy reference datasets^8^. These methods avoid potential biases associated with single-cell clustering and reveal cellular-level phenotypic alterations. In summary, these studies underscore that precise phenotype learning and accurate biological interpretation are crucial for advancing our understanding of diseases.

In this study, we present a deep-learning model named PHenotype prediction with Attention mechanisms for Single-cell Exploring (PHASE). This model utilizes an attention-based neural network framework to predict clinical phenotypes from scRNA-seq data while providing interpretability of key features linked to phenotypic outcomes at both the gene and cell levels. PHASE consists of several components: a data-preprocessing procedure, a gene feature embedding module, a self-attention (SA) module for cell embedding learning, and a pivotal attention-based deep multiple instance learning (AMIL) module for aggregating all single-cell information within a sample. The latent embeddings learned by the model are subsequently used for phenotype prediction. We evaluated PHASE using COVID-19 PBMC scRNA-seq datasets to diagnose infection and predict its severity, covering 560 individuals and over 2.5 million single cells. Additionally, we tested the model on an age-related PBMC single-cell dataset, comprising nearly 2 million single cells, to assess its performance in age prediction and biomarker discovery We demonstrate PHASE outperforms baseline methods in both sample representation learning and phenotype prediction. Notably, by incorporating deep-learning-based interpretability techniques and cell-level attention weights, PHASE reveals interactive alterations across cells and genes, facilitating a more comprehensive understanding of atlas-scale single-cell datasets. As both single-cell atlases and deep learning methodologies continue to evolve, PHASE offers significant advancements in phenotype prediction and provides deeper biological insights from individual cells.

## Results

### The PHASE Framework

To address the potential biases in data integration and cell clustering and to accurately predict clinical phenotypes from single-cell RNA-seq (scRNA-seq) data, thereby understanding phenotype-associated cell states and gene features, we developed a deep-learning model named PHASE (Fig. 1a). This model processes filtered single-cell expression data and predicts phenotypic labels by aggregating information from individual cells while accommodating the variability in single-cell counts across samples. It consists of several key modules: a data-preprocessing step, a gene feature embedding module, a self-attention (SA) module for learning single-cell embeddings, and a pivotal attention-based deep multiple instance learning (AMIL) module for aggregating information across all single cells within a sample^9^ (Fig. 1b). In detail, the model first projects the expression data of highly variable genes (HVGs) from single cells into a low-dimensional embedding space, which is then used for self-attention learning. The SA module captures dependencies between single cells within a sample, allowing the model to learn robust cell embeddings that reflect similar functionalities^10^. Subsequently, the AMIL module applies attention mechanisms to infer the importance (or attention weight) of each cell embedding, combining these weighted embeddings into a comprehensive sample representation. These sample representations are then used in various downstream phenotype prediction tasks, helping to explore the underlying structure of single-cell data in relation to different clinical phenotypes, all within an end-to-end framework.

**Fig. 1.**
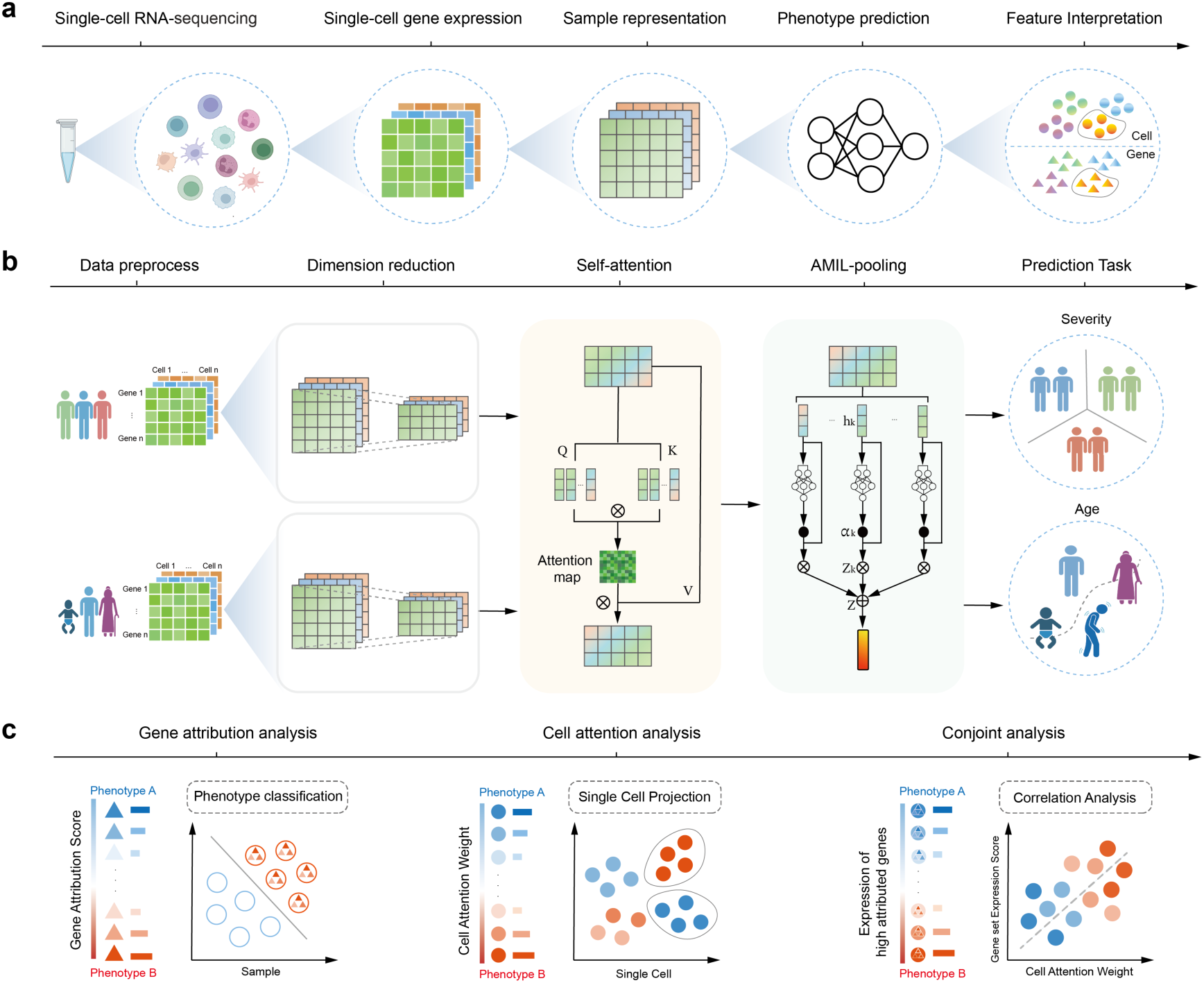
The PHASE Framework. **a**, Schematic diagram illustrating the analysis workflow of the PHASE model. **b**, Diagram of individual modules in the PHASE model pipeline. **c**, Diagram of data interpretation analysis for the PHASE model, featuring gene-level analysis (left), cell-level analysis (middle), and conjoined gene and cell analysis (right).

The interpretability analysis of the PHASE model sheds light on key cell subsets and gene expression features that contribute to clinical phenotype predictions. For gene-level attribution, we employed a deep-learning interpretation method to quantify the contribution of each HVG to phenotype prediction^11^ (Fig. 1c). For cell-level interpretation, we used the inferred cell attention weights to assess the contribution of individual cells to clinical phenotype. Unlike cell frequency, which simply counts the presence of certain cell types, the attention score varies across cell subtypes, indicating that the contribution of cells to phenotype prediction changes depending on their role. We further quantified the relative changes in cellular attention scores in relation to cell frequency, determining whether individual cells contribute more or less to phenotype prediction than their frequency would suggest. Moreover, due to the end-to-end training strategy of our model, we applied the automated cell annotation tool “CellTypist” with predefined cell subset models to uniformly annotate single cells within each sample^12^. This allowed us to summarize cellular attention changes at the cell subset level (Fig. 1c). Since the model simultaneously learns from the full single-cell gene expression matrix, we also conducted joint interpretation analyses of both gene attributions and cell attention weights (Fig. 1c). This approach enables us to accurately identify phenotype-specific single-cell alterations and uncover the underlying biological mechanisms.

### PHASE can predict COVID-19 infection severity

To assess the performance of PHASE in sample classification, we utilized publicly available scRNA-seq data from Peripheral Blood Mononuclear Cell (PBMC) samples of COVID-19-infected patients with distinct disease severity, along with a control cohort of healthy individuals. In total, we obtained 12 public global cohorts, comprising 560 individuals, including 125 healthy samples, 189 mildly infected samples, and 246 severely infected samples^13–24^ (Fig. 2a). The clinical phenotype of each sample was obtained from the original studies and categorized into three groups: H for healthy samples, M for COVID-19 mild/moderate cases, and S for COVID-19 severe/critical cases (Supplementary Table 1). For phenotype classification, we benchmarked PHASE against several deep learning and machine learning methods, including CloudPred^25^, Deep Sets^26^, linear model, and two mixture models (one based on class level and the other on sample level) (Methods). Due to potential biases in sample annotation across these public datasets, we integrated all samples and stratified them into training, validation, and testing sets using a 2:1:1 ratio, conducting 10 independent experiments to ensure robust evaluation. We then evaluated the classification performance using the AUC score, Macro F1-score, precision, and recall metrics, and the result demonstrated that the PHASE model outperforms other models in sample classification (Fig. 2b). For single-cell data interpretation, we trained a final PHASE model using all samples. The Principal Component Analysis (PCA) projection of the learned embeddings of these samples also revealed that the PHASE model was able to clearly distinguish samples based on their phenotype labels (Fig. 2c).

**Fig. 2.**
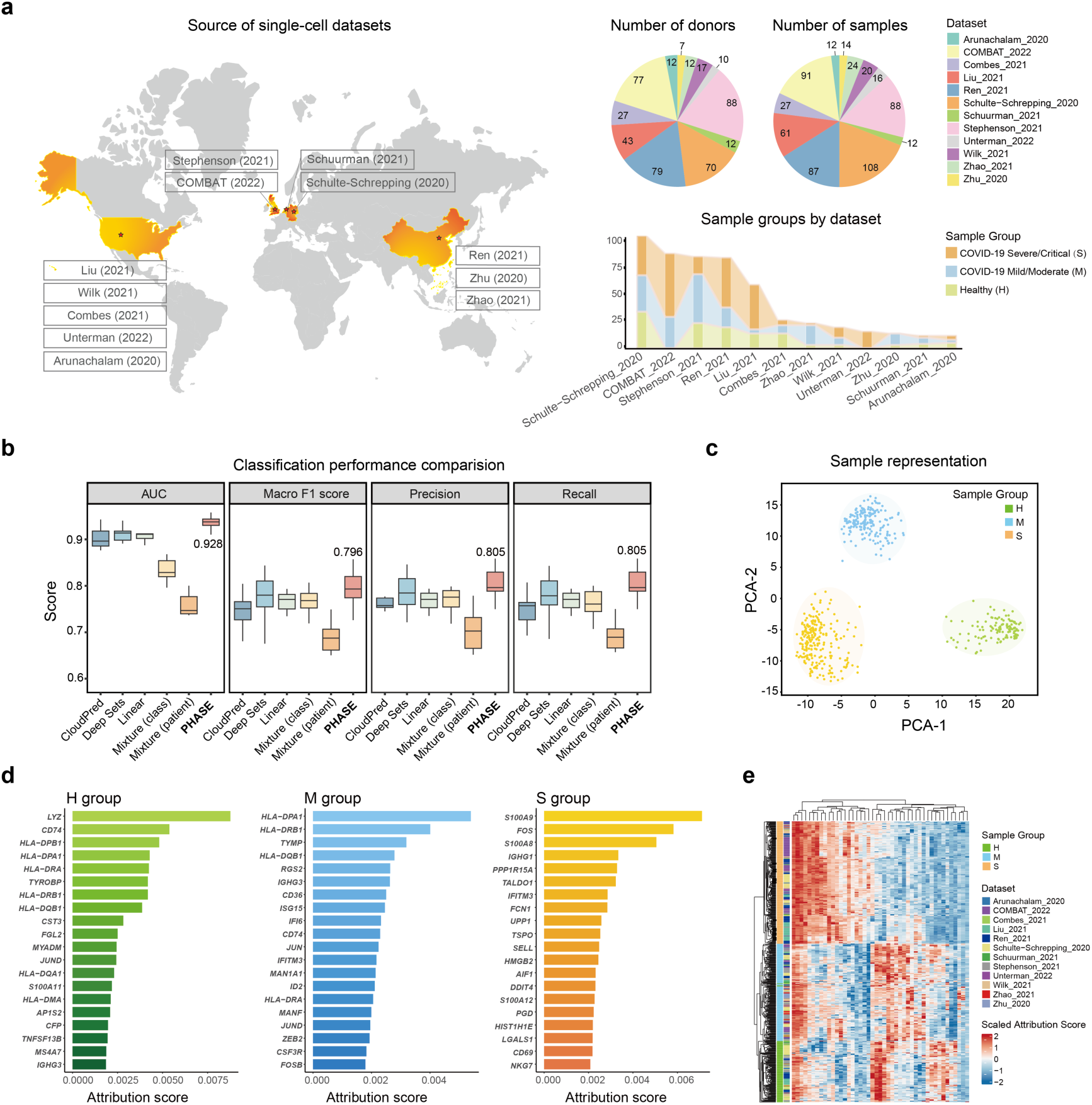
PHASE can predict COVID-19 infection severity. **a**, Overview of the COVID-19 dataset, including sample sources (left), the number of donors, number of samples, and sample group across sub-datasets (right). **b**, Comparison of phenotype classification performance between the PHASE model and other methods. **c,** The PCA projection of sample representation inferred from the PHASE model, colored by the sample group, with confidence ellipses. **d**, Bar plot showing the attribution score of the top-20 high-ranked genes for each sample group. **e**, Heatmap of the scaled attribution score for the selected genes in (d) across individual samples, labeled by sample group and dataset on the left. The heatmap was clustered by hierarchical clustering.

To further explore the unique gene regulatory patterns in different phenotypes and validate the biological insights derived from PHASE, we employed the deep-learning interpretation method Integrated Gradients (IG) to quantify the contribution of various gene features to each phenotype classification^11^. We then ranked the attribution scores of genes associated with each phenotype, highlighting phenotype-specific gene enrichment (Fig. 2d). The high attributed genes in the H group predominantly associated with MHC class II family genes (*CD74*, *HLA-DPB1*, *HLA-DPA1*, *HLA-DRA*, *HLA-DRB1*, *HLA-DQB1*, *HLA-DQA1*, and *HLA-DMA*), while genes in the M group involving interferon-stimulated genes (*ISG15*, *IFI6*, and *IFITM3*), suggesting an antiviral response. In the S group, the release of *S100A8/9* calprotectin has been validated with severely infected COVID-19 patients^27^, as well as the pleiotropic cytokine Galectin-1 (*LGALS1*)^28^ and Pentose Phosphate Pathway (PPP)-related biomarkers *TALDO1* and *PGD*^29^. Moreover, the overlap of highly attributed genes is observed between healthy individuals and those with mild or moderate COVID-19, but not in cases of severe or critical COVID-19. (Suppl. Fig. S1a).

To further confirm the consistency of gene attribution analysis in this study, we compared the average gene attribution values of the top 20 highly attributed genes across all samples and groups, revealing that these genes accurately classified 558 out of 560 samples, independent of their associated gene expression values (Fig. 3e, Suppl. Fig. S1b). This further demonstrates the robustness of the PHASE model in learning single-cell phenotypes.

**Fig. 3.**
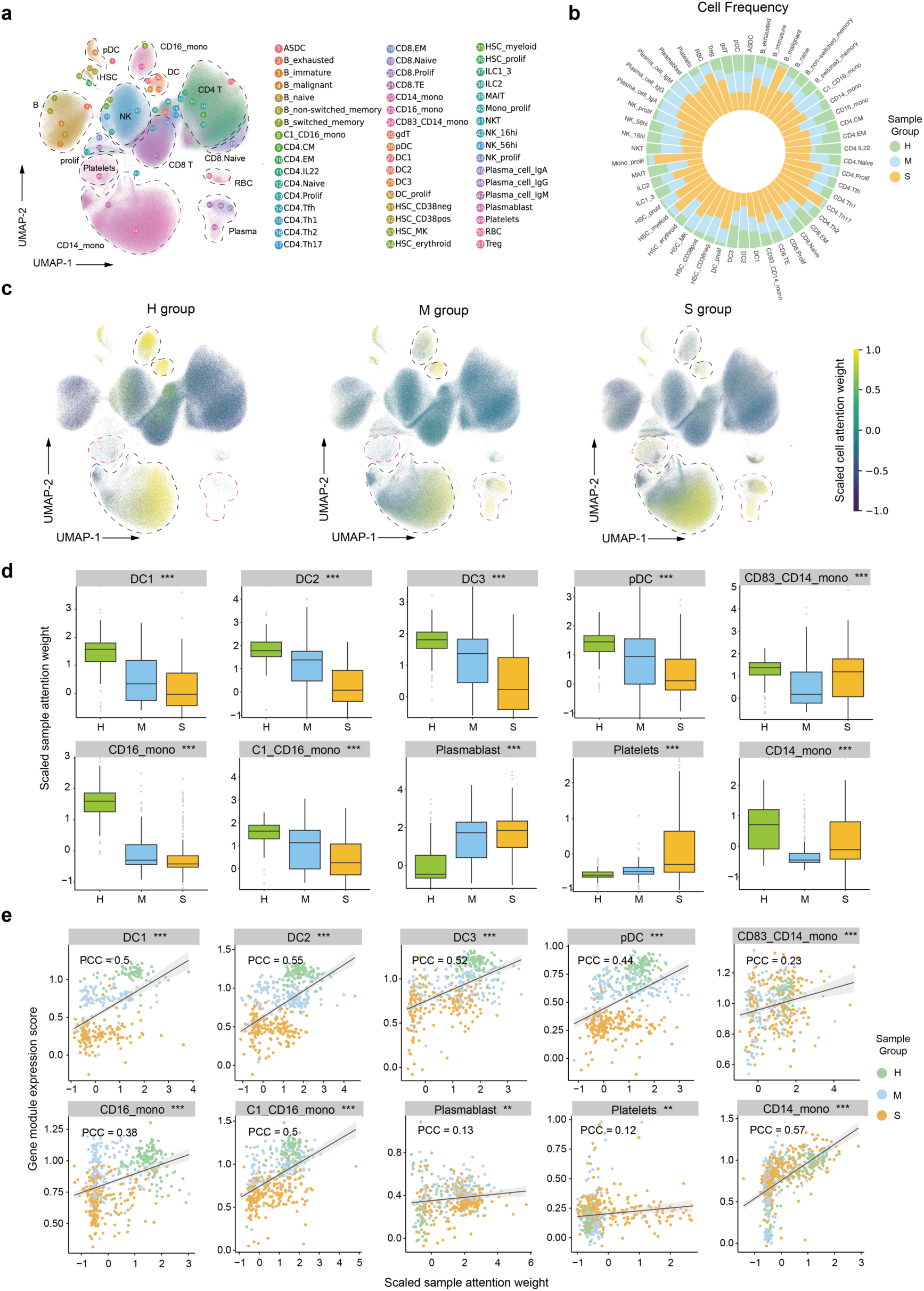
Cellular attention score associated with COVID-19 infection severity. **a**, UMAP plot of single cells from the COVID-19 dataset, colored by automatically annotated cell subsets, and labeled with cell subset ID and major cell subtypes. **b**, The stacked bar plot showing the cell frequency of sample groups across individual cell subsets. **c**, UMAP plots of single-cells as in (a), spitted by sample groups, and colored by scaled cell attention weight. **d**, Comparison of scaled sample attention weights within individual cell subsets across sample groups. Boxplots represent the median, upper quartile, and lower quartile values for each group. **e**, Pearson’s correlation analysis between the sample gene module expressions of group-specific high attributed genes and the scaled sample attention weights. The plot is labeled with correlation coefficient, a regression line, and its confidence interval. The Kruskal-Wallis test was used to assess statistical significance in (d). The statistical levels indicated as follows: *p ≤ 0.05, **p ≤ 0.01, ***p ≤ 0.001 in (d and e).

### Cellular attention score associated with COVID-19 infection severity

To summarize the cell-level association with clinical phenotypes, we obtained the inferred attention score of each cell within a sample. Similar to cell frequency, the sum of attention scores in each sample equals one. However, due to varying cell counts across samples, raw attention scores are not directly comparable between samples. To address this, we used the log2-transformed ratio of the cellular attention score to its average cell frequency, providing a measure of the relative increase or decrease in attention for individual cells within a sample (Methods). To further aggregate the relative attention scores at the cell subset level, we employed the CellTypist method and the “Healthy_COVID19_PBMC” model to annotate single-cell subtypes in each sample^30^ (Supplementary Table 2). The median relative attention scores of single cells were used to represent the cell subset-level score in a sample. Additionally, z-score-scaled relative attention scores at both the single-cell and cell subset levels were used for downstream analyses.

To identify the COVID-19 infection-related single-cell subsets, we integrated all single cells into a Uniform Manifold Approximation and Projection (UMAP) map, labeling them based on automatically annotated single-cell subsets (Fig. 3a). By comparing the distributions of these subsets across phenotype groups and their relative attention scores (Fig. 3b, c), we identified the cell-type-specific enrichment in attention scores. This suggests that the AMIL module successfully uncovers the distinct cellular contributions to different clinical phenotypes. We then analyzed the relative attention scores of different cell subsets across phenotype groups, identifying an enrichment of DC1-3 cells, pDC cells, and CD16^+^ monocytes in the healthy cohort. In contrast, we observed an attention enrichment of CD14^+^ monocytes and platelets in severely infected patients, as well as the enrichment of plasmablast cells in all COVID-19 infected patients (Fig. 3d and Suppl. Fig. S2). A significant reduction of dendritic cells has been observed in COVID-19 patients^29^. The shift across distinct monocyte subsets and the increased attention weight of platelets suggest the alternated innate immune response and potential platelet-monocyte communication related to disease severity^31,32^.

For assessing the potential consistency across single-cell attention scores and group-specific high-attribute genes, we calculated the Pearson’s correlation coefficients (PCC) between relative attention weight and gene expression scores of group-specific top-50 high-attributed genes in each group. The results demonstrated strong correlations between attention scores and gene module scores in dendritic cells, CD14^+^ and CD16^+^ monocytes (Fig. 3e, Suppl. Fig. S3). This finding aligns with the observed reduction in the expression of HLA-DR family genes in severe COVID-19 cases^33^, suggesting that the PHASE model effectively enriches both highly contributing single cells and their corresponding gene expression patterns for phenotype prediction.

### PHASE can predict age with PBMC single-cell datasets

We then assess the performance of the PHASE model in regression learning, predicting the age label of a PBMC single-cell dataset of healthy people, and uncovering the age-related single-cell profiles. In detail, scRNA-seq data from 317 samples were obtained from the study of Terekhova et al.^34^, as well as the age label, age group, and cellular annotations (Fig. 4a, Supplementary Table 3). The age of this cohort ranges from 25 to 85 years old, we then split this dataset into training and validation datasets with a 4:1 ratio and performed 5-fold cross-validation. As described before, the sample representations aggregated from the AMIL module were used for age regression prediction. The PCC of predicted age with true age labels for each fold ranges from 0.82-0.89, demonstrating the PHASE model can successfully learn the age-related single-cell features (Fig. 4b). The result is also better than the baseline linear prediction model (Fig. 4c). Additionally, to validate the performance of PHASE model in age prediction, we utilized another testing dataset from Zhu et al.^35^, which include 17 samples range from 28 to 77 years (Supplementary Table 3). Besides, limited to the data bias across different studies, 4265 of 5000 (85.3%) HVGs in the training dataset were used for model testing, and the expression of absent genes was set to zero. The PCC of the testing dataset was 0.64, further proving the performance of the PHASE model in age prediction (Fig. 4d).

**Fig. 4.**
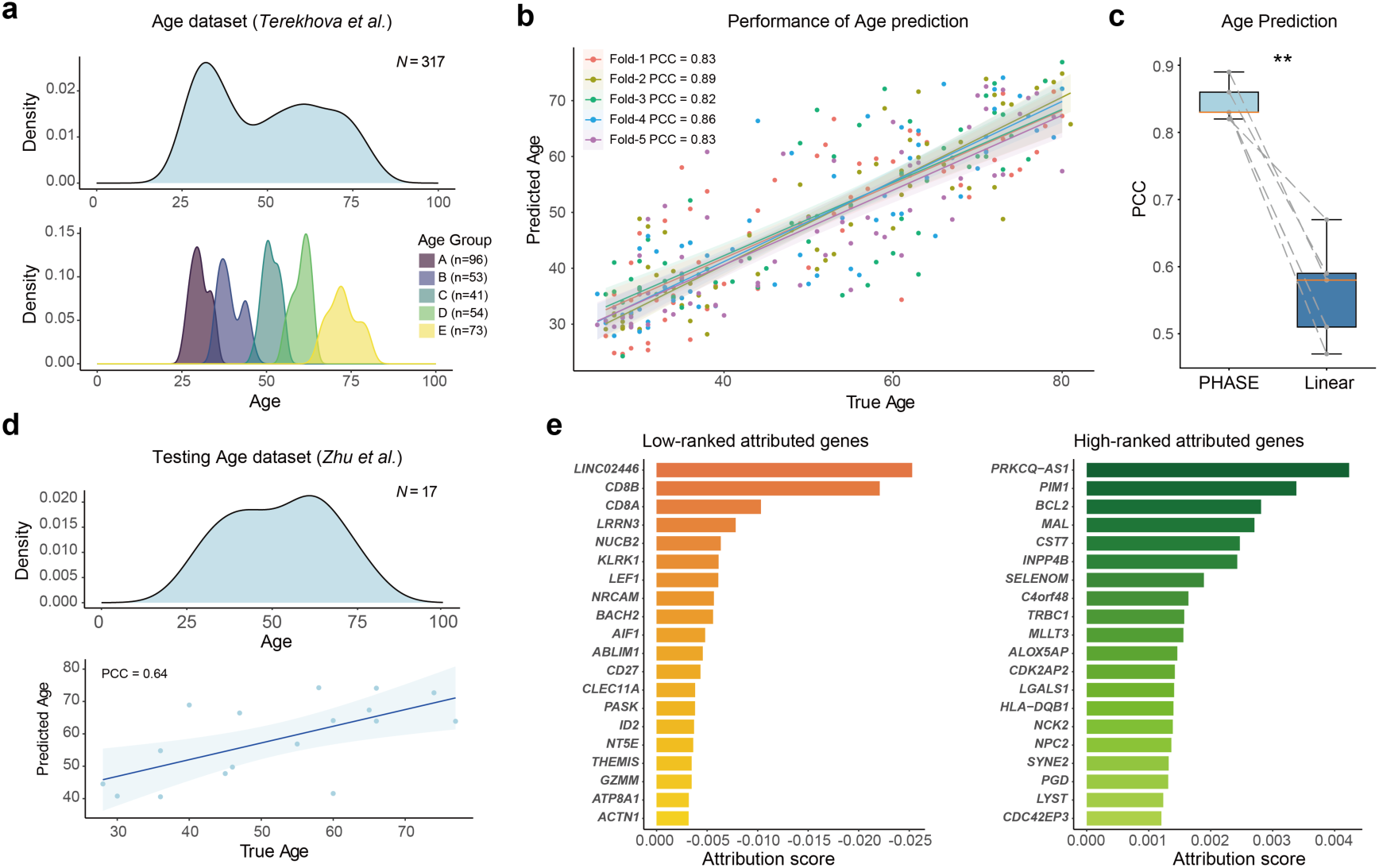
PHASE can predict age with PBMC single-cell datasets. **a**, Overview of the age distribution in the Age_Terekhova_2023 dataset, displayed by individual samples (top) and by age groups (bottom). **b,** Scatter plot showing the true versus predicted age for the PHASE model using test data from 5-fold cross-validation, colored by each fold. The plot is labeled with Pearson’s correlation coefficient (PCC), a regression line, and its confidence interval for each fold. **c,** Comparison of age prediction performance between the PHASE model and a Linear method. The paired PCC for the test data in each fold, as shown in (b), was used for comparison. **d,** Overview of the age distribution in the Age_Zhu_2023 dataset, displayed by individual samples (top), and scatter plot showing the true versus predicted age for the PHASE model with this dataset. The plot is labeled with the PCC, a regression line, and its confidence interval. **e,** Bar plot showing the attribution score of the 20 low-ranked (left) and high-ranked (right) genes in age prediction of the PHASE model. The paired Kruskal-Wallis test was used to assess statistical significance in (c), with *** indicated p ≤ 0.001.

Similar to deep-learning model interpretation analysis in the classification task, we identified the contribution of gene features in age prediction, and the positive IG values mean highly contribution of elder samples and, *vise vase*. Thereby, the high-ranked and low-ranked genes indicated the age-related gene features (Fig. 4d). The expression of leucine-rich repeat neuronal 3 (*LRRN3*) has been observed to decrease in elder samples^36^, as well as the age-related immunosenescence (*LEF1*)^37^ and autophagy disruption (*BCL2*)^38^.

### Cell attention for interpreting age-related single-cell features

**We then assessed the relationship between cell attention weights of single-cell subsets and their age labels, and** used the same cellular annotations as in Terekhova et al.^34^ **for** direct comparison (Suppl. Fig. S4). By comparing z-score-scaled relative attention scores across different age groups, we observed a distinct age-specific enrichment of cellular attention across various cell subsets (Fig. 5a). This suggests that the PHASE model effectively captures age-related single-cell features. Similar to cell frequency, the sum of cell attention scores reflects the contribution of specific cell subsets to phenotype prediction, facilitating a paired comparison between cell frequency and attention score. To quantify the alternation between cell frequency and cell attention score, we calculate the Root Mean Square Error (RMSE) metric with each other based on major cell subsets (Methods). Notably, we found that the RMSE values for individual samples decreased with advancing age (Fig. 5b), suggesting that the PHASE model adjusts the attention values of single cells in younger samples while anchoring elder samples. Moreover, we estimated the RMSE metric across major cell populations (Methods) and observed that the myeloid cell, TRAV1-2^-^ CD8^+^ T cells, and CD4^+^ T cells were mostly affected in cell attention scores, suggesting their potential roles in age prediction (Fig. 5c).

**Fig. 5.**
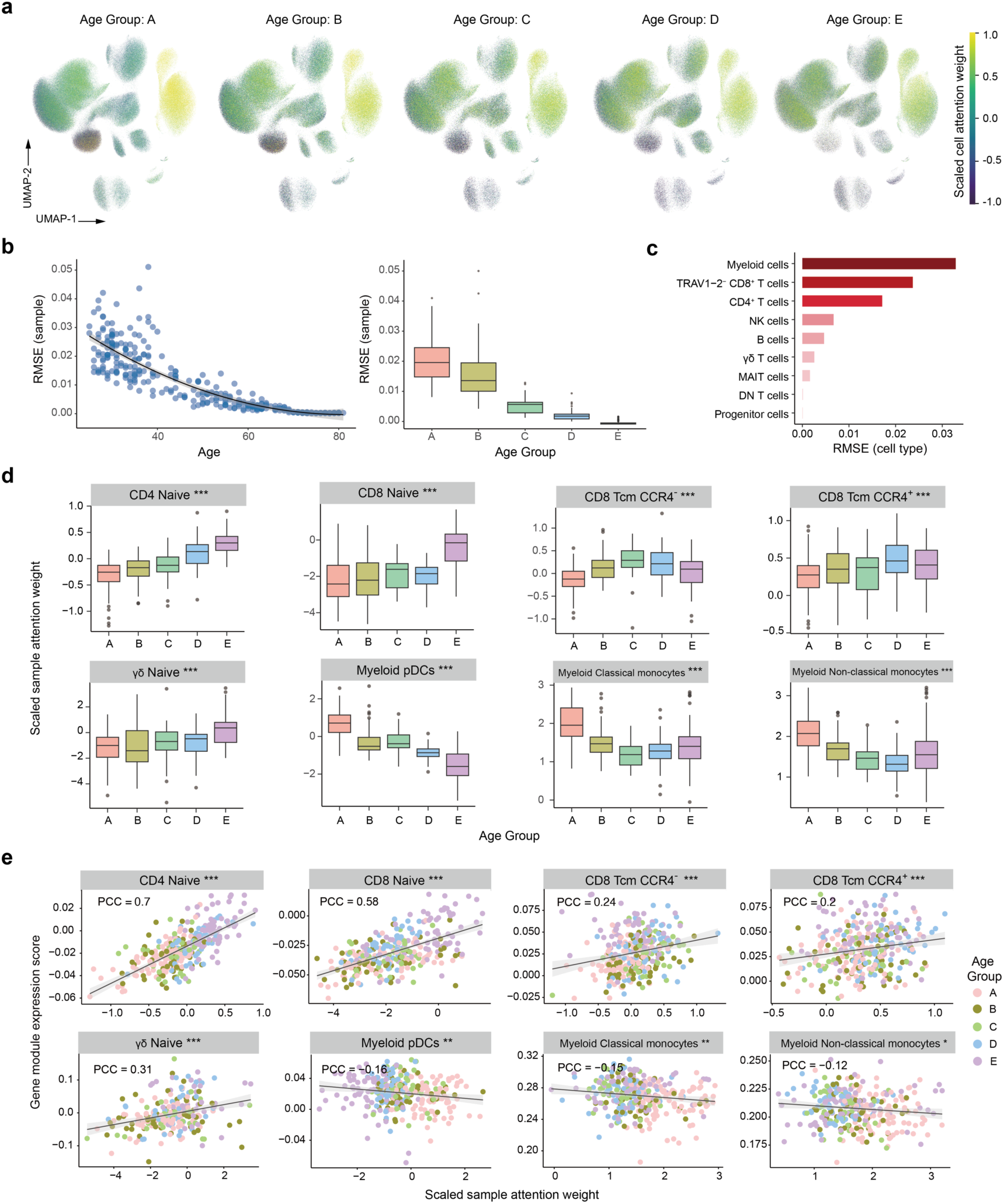
Cell attention for interpreting age-related single-cell features. **a**, UMAP plots of single-cells in the Age_Terekhova_2023 dataset, split by age groups and colored by scaled cell attention weight. **b,** Distribution of calculated Root Mean Square Error (RMSE) between cell frequency and attention scores for each sample across different ages (left) and age groups (right). The bivariate regression line and its confidence interval were labeled on the left panel. **c,** Distribution of calculated Root Mean Square Error (RMSE) between cell frequency and attention scores of all samples, group by major cell type. **d,** Comparison of scaled sample attention weights within individual cell subsets across age groups. Boxplots represent the median, upper quartile, and lower quartile values for each group. **e**, Pearson’s correlation analysis between the sample gene module expressions of age-related high-ranked attributed genes and the scaled sample attention weights. The plot is labeled with a correlation coefficient, a regression line, and its confidence interval. The Kruskal-Wallis test was used to assess statistical significance in (d). The statistical levels indicated as follows: *p ≤ 0.05, **p ≤ 0.01, ***p ≤ 0.001 in (d and e).

We further analyzed the contributions of specific cell subsets to age prediction across major cell populations. **Unlike the classification task, we utilized the coefficient of determination (R^2^) metric to quantify the relationship between z-score-scaled relative attention scores and age labels (Methods)**. We ranked the R^2^ scores for cell subsets within each major cell population (Suppl. Fig. S5), and found that most highly-ranked cell subsets were defined as age-related in Terekhova et al.^34^. While the frequency of naïve cells declined in older samples, the relative attention scores for naïve CD4^+^, CD8^+^, and ψο T cells increased, suggesting that the functional and abundant changes of these cells are significant for age prediction, along with the *CCR4*^+/-^ Tcm CD8^+^ T cells (Fig. 5d). Moreover, despite the relatively stable frequency of myeloid cells noted by Terekhova et al.^34^, we observed that the relative attention scores of most myeloid cells, including pDCs, classical and non-classical monocytes decreased with age, suggesting their potential roles in predicting younger samples (Fig. 5d, Suppl. Fig. S6). As described before, we evaluated the gene module expression of top-100 most highly age-attributed genes with relative cell attention weight, to uncover the biological mechanisms linking attention scores to age (Fig. 5e). Notably, we found a positive correlation of these indexes in T cells, particularly in naïve CD4^+^ and CD8^+^ T cells, but negative in myeloid cell subsets (Fig. 5e, Suppl. Fig. S7). These results further validate that both gene attribution and cell attention scores in the PHASE model reflect age-related changes in single-cell datasets, and also propose that phenotype changes in naïve T cells can be a valuable biomarker in age prediction.

## Discussion

Single-cell RNA sequencing (scRNA-seq) has emerged as a critical tool for understanding cellular heterogeneity and its relation to disease phenotypes^1^. Recognizing the cell subpopulations and genes associated with disease phenotype from single-cell data is of fundamental importance for developing cell population-specific targeted therapies and the discovery of biological biomarkers^39,40^. In this study, we present PHASE, a novel deep learning framework designed to integrate single-cell RNA sequencing (scRNA-seq) data for the prediction of clinical phenotypes. The model addresses several key challenges in single-cell analysis, including data integration biases, variable cell counts across samples, and the complexity of aggregating cellular features to uncover phenotype-specific alterations. Our results demonstrate the utility of PHASE in predicting clinical outcomes, such as COVID-19 infection severity and age, while also advancing the interpretability of single-cell data through attention mechanisms.

The challenges in interpreting these high-dimensional single-cell datasets persist due to the variability in cell populations and gene expression across samples. PHASE addresses these challenges by implementing a self-attention (SA) module to capture dependencies between single cells within a sample and an attention-based multiple instance learning (AMIL) module to aggregate this information across the entire dataset. The model’s ability to learn robust cell embeddings and attribute importance to specific cells and genes greatly enhances the biological interpretability of scRNA-seq data, distinguishing it from other single-cell phenotype prediction approaches^25,26^. The end-to-end framework of our model differs from an integration-based single-cell multimodal MIL prediction model^41^ but is similar to the whole-slide image-based phenotype prediction approach^42^. By aggregating individual cell features into a unified sample representation, PHASE provides a new approach to uncovering meaningful insights into the relationship between cellular states and disease phenotypes.

One of the key contributions of PHASE is its ability to enhance the interpretability of single-cell data using attention mechanisms, which advances beyond traditional cell-type frequency analysis, offering a deeper understanding of how individual cells contribute to clinical outcomes.

This is an important advancement because it allows for the identification of both highly abundant and rare cell types that are crucial for understanding disease mechanisms. For instance, in the case of COVID-19, our results showed that certain CD14^+^ monocyte populations, platelets, and plasmablasts had enriched attention scores in severe cases, while dendritic cells (DCs) were more prominent in healthy individuals. This cell-specific attention distribution suggests that PHASE can accurately capture the functional importance of different immune cell subsets in response to infection severity. PHASE also performed robustly in predicting age-related changes in single-cell profiles, demonstrating the model’s flexibility in handling both classification and regression tasks. The attention mechanisms enabled us to detect subtle changes in cellular attention, particularly in naïve T cells, which were identified as key contributors to age prediction. Furthermore, the consistency between high attributed genes and attention weight of specific cell populations, such as dendritic cells and monocytes, further validates PHASE’s capacity to integrate both gene-level and cell-level contributions to phenotype prediction.

Despite the promising results, there are several limitations to the current study that warrant further investigation. First, while PHASE demonstrated strong performance in predicting COVID-19 severity and age from scRNA-seq data, its generalizability to other diseases and phenotypes remains to be fully explored. The PHASE model also relied on an atlas-level single-cell dataset to uncover robust phenotype-specific single-cell alternations, limiting its application to small-size cohorts. The model’s reliance on pre-defined cell annotation models, such as CellTypist, introduces potential biases in cell subtype identification, particularly when applied to datasets from different biological contexts or conditions not well-represented in the training data. Moreover, while attention mechanisms improve the interpretability of deep learning models, they can also introduce challenges in distinguishing between biologically meaningful patterns and artifacts arising from the model architecture. Further validation through experimental methods, such as single-cell perturbation studies, will be necessary to confirm the biological relevance of the cells and genes identified by PHASE as important contributors to disease phenotypes.

In conclusion, PHASE offers significant advancements in single-cell RNA-seq analysis by integrating attention mechanisms to predict clinical phenotypes while providing interpretable insights into cell and gene contributions. The model’s performance in predicting COVID-19 infection severity and age highlights its potential for broader applications in disease research. However, future work should focus on improving the model’s generalizability and validating its predictions. These efforts will further enhance our ability to leverage single-cell data for precision medicine and biomarker discovery.

## Methods

### Data collection and preprocessing COVID-19 scRNA-seq data collection

We sourced raw gene expression counts from a compendium of 560 COVID-19 PBMC 10X Genomics scRNA-seq datasets across 12 studies. The clinical phenotype for each sample was determined in the original studies, which categorized patients into mild, moderate, severe, or critical COVID-19 conditions, following the WHO ordinal scale for disease severity: https://www.who.int/westernpacific/emergencies/covid-19/information/asymptomatic-covid-19. For convenience, we categorized all samples into three groups: H (healthy), M (mild and moderate), and S (severe and critical). Our collection comprised 246 samples from patients with severe/critical COVID-19 (S), 189 samples from those with mild/moderate symptoms (M), and 125 samples from healthy controls (H).

**COVID-19_Arunachalam_2020.** We obtained the COVID-19 PBMC dataset from ref. 13, which contains 12 samples and 49,847 cells, comprising 4 samples from the S group (severe/critical patients), 3 from the M group (mild/moderate patients), and 5 from the H group (healthy controls).

**COVID-19_COMBAT_2022.** We obtained the COVID-19 PBMC dataset from ref. 14, which contains 91 samples and 560,546 cells, comprising 61 samples from the S group and 30 from the H group.

**COVID-19_Combes_2021.** We obtained the COVID-19 PBMC dataset from ref. 15, which contains 27 samples and 49,672 cells, comprising 5 samples from the S group, 8 from the M group, and 14 from the H group.

**COVID-19_Liu_2021.** We obtained the COVID-19 PBMC dataset from ref. 16, which contains 61 samples and 346,542 cells, comprising 43 samples from the S group, 4 from the M group, and 14 from the H group

**COVID-19_Ren_2021.** We obtained the COVID-19 PBMC dataset from ref. 17, which contains 87 samples and 512,163 cells, comprising 48 samples from the S group, 19 from the M group, and 20 from the H group.

**COVID-19_Schulte-Schrepping_2020.** We obtained the COVID-19 PBMC dataset from ref. 18, which contains 108 samples and 211,056 cells, comprising 38 samples from the S group, 35 from the M group, and 35 from the H group.

**COVID-19_Schuurman_2021.** We obtained the COVID-19 PBMC dataset from ref. 19, which contains 12 samples and 8,460 cells, comprising 2 samples from the S group, 6 from the M group, and 4 from the H group.

**COVID-19_Stephenson_2021.** We obtained the COVID-19 PBMC dataset from ref. 20, which contains 88 samples and 486,419 cells, comprising 17 samples from the S group 47 from the M group, and 24 from the H group.

**COVID-19_Unterman_2022.** We obtained the COVID-19 PBMC dataset from ref. 21, which contains 16 samples and 66,223 cells, with all samples coming from the S group.

**COVID-19_Wilk_2021.** We obtained the COVID-19 PBMC dataset from ref. 29, which contains 20 samples and 165,700 cells, comprising 10 samples from the S group 7 from the M group, and 3 from the H group.

**COVID-19_Zhao_2021.** We obtained the COVID-19 PBMC dataset from ref. 22, which contains 24 samples and 54,912 cells, comprising 17 samples from the S group 47 from the M group, and 24 from the H group.

**COVID-19_Zhu_2020.** We obtained the COVID-19 PBMC dataset from ref. 23, which contains 14 samples and 29,089 cells, comprising 11 samples from the M group, and 3 from the H group.

### COVID-19 scRNA-seq data preprocessing

We employed the Scanpy toolkit^43^ within the Python platform to manage and analyze all single-cell data. In the initial quality control process, cells with fewer than 200 detected mRNA molecules and exhibiting over 10% mitochondrial gene expression were excluded, and genes expressed in fewer than 3 cells were filtered out. This process yielded a refined dataset of 2,540,629 high-quality cells, with an average of 4,537 cells per sample. Following quality control, we normalized the expression matrix and applied a logarithmic plus one transformation to prepare the data for downstream analysis. Using Scanpy’s ‘sc.pp.highly_variable_genes’ function and using sample name as the batch label, we identified 5,000 highly variable genes based on the dispersion of log-normalized counts. To ensure consistent cell type annotation, we utilized the “Healthy_COVID19_PBMC.pkl” model from the CellTypist toolkit^12,44^ (https://www.celltypist.org/models), leading to the identification of 51 distinct cell subtypes across the 2,540,629 cells. The finalized dataset, including the annotated cells and highly variable genes, was saved in h5ad format for subsequent analysis.

### Age scRNA-seq data collection

We obtained Age_Terekhova_2023 from the Synapse database with the code <u>syn49637038</u>, which provided preprocessed data stored in h5ad files. The raw data of Age_Zhu_2023 can be accessed on the GEO database with the code <u>GSE213516</u>.

**Age_Terekhova_2023.** We obtained the age PBMC dataset from ref. 34, which contains 317 samples and 1,916,367 cells ranging from 25 to 85 years old.

**Age_Zhu_2023.** We obtained the age PBMC dataset from ref. 35, which contains 17 samples and 132,954 cells ranging from 28 to 77 years old.

### Age scRNA-seq data preprocessing

For preprocessed data stored in h5ad files from the Age_Terekhova_2023 dataset, we identified 5,000 highly variable genes based on the dispersion of log-normalized counts, as described before. Cell type annotations were adopted directly from the original publication of the raw data. The finalized dataset, which includes the annotated cells and the expression of highly variable genes, was saved in h5ad format for subsequent analysis. For raw data from the Age_Zhu_2023 dataset, we used the ‘sc.read_10x_mtx’ function from the Scanpy toolkit to convert the data into AnnData format, filtering out cells with fewer than 200 detected genes or more than 10% mitochondrial gene expression. We normalized the expression matrix, applied a logarithmic transformation, and then subset the expression data to include only the intersection of genes between Age_Zhu_2023 and the highly variable genes (HVGs) from Age_Terekhova_2023, filling in missing genes with zeros. The processed dataset was saved as a h5ad file for further analysis.

### PHASE model framework

PHASE is an algorithm designed to predict clinical phenotypes from single-cell RNA-seq data by integrating association in cell states and gene features. It takes a single-cell expression matrix, filtered for highly variable genes as input, and predicts phenotypic classification results as output. The algorithm consists of several key components: a data preprocessing module, a gene feature embedding module, a self-attention (SA) module for learning cell similarities, and an attention-based deep multiple instance learning (AMIL) module, which aggregates cell data and evaluates the contribution of each cell to phenotype prediction.

### Data-preprocessing procedure

We employed the ‘Scanpy’ and ‘AnnData’ packages to load the finalized dataset saved in h5ad format. Iterating through each sample identifier in the dataset, we extracted the corresponding single-cell data and converted the expression matrices into floating-point tensors. For the classification task, samples were grouped into three categories based on the ‘group’ attribute in the AnnData subject: healthy (0), mild/moderate COVID-19 infection (1), and severe/critical COVID-19 infection (2). Labels were encoded using scikit-learn’s ‘LabelEncoder’ function, and both the data tensors and label tensors were serialized with the pickle module. For the regression task, the gene expression matrix was converted to a tensor for each sample, with the corresponding age attribute serving as the regression label. The serialized data was saved as pickle files in designated directories to facilitate subsequent model training and analysis.

### Gene feature embedding module

This module is responsible for the initial embedding of gene features. It utilizes a fully connected layer to reduce the dimensionality of the input from 5,000 gene features down to 256 dimensions. After this reduction, a ReLU activation function is applied to enhance the model’s non-linear processing capabilities, and a dropout layer with a rate of 0.25 is used to mitigate overfitting. The output of this module is the embedding feature representation of each single cell, which is then passed to the subsequent self-attention processing stage.

### Self-attention (SA) module

In this study, the self-attention mechanism leverages a Transformer encoder structure was used to process single-cell RNA-seq data. Each cell is represented as a 256-dimensional vector within a matrix of dimensions n × 256, where n is the number of cells. The attention mechanism functions as follows:

Input feature vectors are transformed into query (Q), key (K), and value (V) matrices by multiplying the input matrix with trainable weight matrices specific to Q, K, and V. The Attention scores are calculated using the formula:

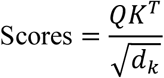

Where *d_k_* is the dimension of the key vectors. This step measures the influence of each cell on every other cell, with the dot product scaled down to stabilize gradients.

Next, the scores are normalized across each row using the SoftMax function to generate an attention map, dictating the relevance of each cell’s features relative to others:

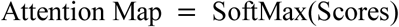

The final output feature vector for each cell is computed as a weighted sum of the value vectors, updated by the attention weights:

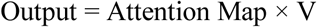

This results in a n × 256 matrix, where each column represents an updated feature vector for each cell.

The multi-head attention is introduced to enhance the model’s capacity to focus on different parts of the input data simultaneously. In multi-head attention, multiple attention heads are used, each with its own set of Q, K, and V weight matrices. The attention mechanism is applied in parallel across each head, allowing the model to capture diverse patterns and relationships within the data. The outputs of the individual heads are concatenated and linearly transformed to produce the final output:

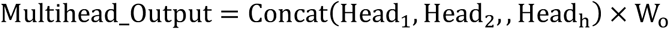

where *h* is the number of attention heads, and *W*_D_ is the output projection matrix. This enables the model to attend to different aspects of the cell interactions in parallel, further improving performance and representation learning of the data.

### Attention-based deep multiple instance learning (AMIL) module

In this model, the Attention-Based Deep Multiple Instance Learning (AMIL) mechanism^9^ is implemented to effectively aggregate the feature representations of individual cells within a sample, enabling a comprehensive evaluation of related disease phenotypes. Each cell in the dataset is represented by an instance feature vector *h*_k_, which encapsulates the gene expression profile specific to that cell. The relevance of each cell in representing the disease phenotype is quantitatively assessed through a gating and attention mechanism, which computes weights *α_k_*, for each cell. The gating mechanism utilizes a combination of hyperbolic tangent (tanh) and sigmoid (σ) functions to the transformed feature vectors:

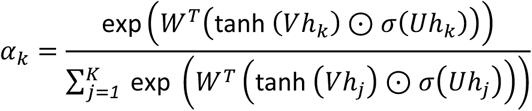

Here, *V* and *U* are trainable weight matrices that modulate the feature vectors through tanh and sigmoid activations, respectively. This allows the model to focus on relevant features while suppressing less informative ones. The weights *α_k_*represent the importance of each cell’s contribution to the disease phenotype, derived by normalizing the exponential scores across all cells to ensure they sum to one.

The final representation of the sample *Z*, is then calculated as a weighted sum of all instance feature vectors, reflecting the aggregate influence of all cells:

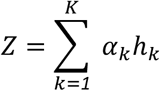

This aggregated feature vector *Z*, is used to predict the overall disease phenotype, effectively capturing the collective impact of individual cellular profiles on the disease manifestation.

### PHASE training

For the classification task, PHASE was optimized using Adam the Adam optimizer with a learning rate of 0.00001. CrossEntropyLoss was employed as the loss function, and the model was trained for 300 epochs. Similarly, for the regression task, PHASE was optimized using Adam with a learning rate of 0.000005, and MSELoss as the loss function. In this case, the model was trained for 1000 epochs. For both tasks, after model evaluation, we trained a final model using all datasets to enable precise single-cell interpretation of both cellular and gene features. All experimental procedures were conducted on a workstation equipped with AMD EPYC 9754 128-core Processor CPUs and NVIDIA GeForce RTX 4090 GPUs.

### Benchmarking methods

We conducted a benchmark to compare PHASE with five other phenotype prediction methods: CloudPred^25^, Deep Sets^26^, linear model, and two mixture models (one based on class level and the other on sample level). The data-splitting strategy for the benchmarking methods follows the PHASE model, and the final result is determined by averaging the performance across all iterations. We evaluated the performance of these methods on our finalized COVID-19 and age datasets. **We followed the instructions provided on the CloudPred GitHub repository:** https://github.com/bryanhe/CloudPred. The detailed configurations for each method are as follows:

CloudPred For our implementation, we set the parameters to ‘iterations = 500’ and ‘lr = 1e-4’. The model employs a ‘DensityClassifier’ to probabilistically model the clusters using Gaussian distributions, allowing it to predict disease phenotypes by identifying and quantifying the contributions of various cellular subpopulations, and we set the classifier parameter with ‘state = 3’.

**Deep_Sets The predicted phenotype was determined using an ‘Aggregator’ function, which maintains invariance by combining features regardless of the order of elements. We set the parameters to ‘iterations = 1000’ and ‘lr = 1e-4’.**

**Linear** Linear processes single-cell RNA-seq data by treating each cell as an independent entity. It employs a single linear layer to map input dimensions to outputs, predicting phenotypes for individual cells. **We use the parameter ‘iterations = 400, lr = 1e-4’. Additionally, we applied this approach to predict age labels using a regression prediction network.**

### Mixture (Class)

The predicted phenotype using a ‘GaussianMixture’ function to fit the single-cell data of each group.

### Mixture (Patient)

The predicted phenotype using a ‘GaussianMixture’ function to fit the single-cell data of each sample.

## Metrics

### Gene attribution analysis associated with phenotype

We employ the Integrated Gradients (IG) method to calculate gene attribution scores related to different clinical phenotypes. This method estimates the attribution of each gene by integrating the gradients along the path from a baseline (a zero vector) to the actual input, quantitatively expressing how each gene’s expression levels contribute to the model’s predictions. The formula used is as follows:

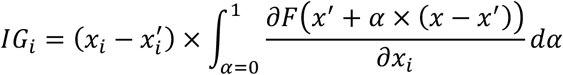

where *x* represents the input, *x*^’^ is the baseline, α scales the input from the baseline to the actual input, 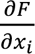 and represents the gradient of the model’s output with respect to the input feature *x*_Z_.

For the classification task, we computed the sample-attribution score for each gene across individual samples. To obtain the group-attribution score for each gene, we averaged the sample-attribution scores within each group. For the regression task, we computed the sample-attribution score for each gene across individual samples. The gene-attribution score was then calculated by averaging the sample-attribution scores for each gene across all samples.

### Estimation of Cellular Attention Weights Associated with Phenotypes

We employ attention-based deep multiple instance learning (AMIL) to calculate cell attention scores associated with COVID-19 severity or age. The cell-attention scores α’ represent the importance of each cell within a sample, as previously described. To ensure the comparability of these scores across different samples, we applied a z-score scaling transformation to the cell-level attention scores. Specifically, for each cell *k*, the attention score α_k_ was scaled using the following formula:

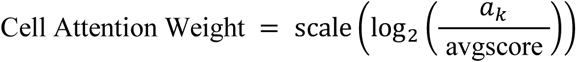

Where the “avgscore” represents the average importance score for all cells within a sample, computed as:

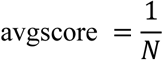

with *N* being the total number of cells in the sample.

To compute cell type-level attention weights, we first calculated the median of the log-transformed attention scores across all cells of the same type, followed by z-score scaling. This process is described by the following formula:

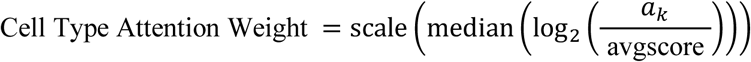

For both classification and regression tasks, we calculated the attention scores at both the cell and cell type levels for individual samples. This method ensures normalized and comparable assessments of cellular and cell type importance across different samples.

### RMSE calculation

We used Root Mean Square Error (RMSE) to quantify the deviation between cell attention scores and cell frequencies, a lower RMSE values indicate better alignment between these two metrics. For sample-level RMSE calculation, we calculated the RMSE between the sum of cell type attention scores for each cell type and the corresponding cell frequencies. The RMSE for each sample is defined as:

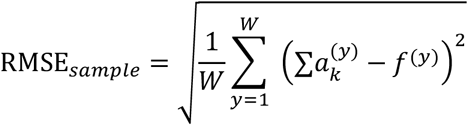

Where *W* represents the number of cell types, *a*^(y)^_k_ represents the summed cell attention score for cell type *y*, and *f*^(z)^ denotes the corresponding cell frequency for cell type *y*.

For cell-type-level RMSE calculation, we calculated the RMSE between the sum of cell type attention scores and the corresponding cell frequencies across all samples. The RMSE for each cell type is defined as:

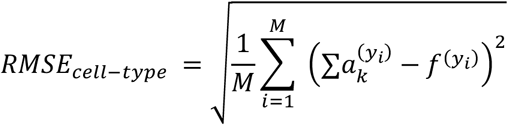

Where *M* represents the number of samples, *a*^(yi)^_k_ represents the summed attention scores for cell type *y*_i_ in sample *i, f*^(yi)^ denotes the corresponding cell frequency for cell type *y* in sample *i*.

### R^2^ calculation

The R^2^ (coefficient of determination) statistic was used to quantify the degree of correlation between cell attention scores and age, where higher R^2^ values indicate a stronger linear relationship between these two factors. For each cell type, we calculated R^2^, which assesses the proportion of variance in the dependent variable (age) that is explained by the independent variable (cell attention scores). The R^2^ for each cell type is defined as follows:

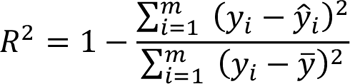

Where *f*_Z_ represents the actual observed value of the dependent variable (age) for sample *i*, *y*^^^_i_ is the predicted value of age based on the independent variable (cell attention scores) for sample *i*, *y*^-^ is the mean of observed values of ages, and *m* is the number of samples.

### Correlation estimation across cell attention and gene module expression

We calculated the gene expression score for each cell using the ‘scanpy.tl.score_genes’ function, which quantifies the expression levels of the selected gene modules to random control genes. We then performed Pearson’s correlation analysis to assess the correlation between the calculated gene expression scores and the corresponding cell attention scores.

## Code availability

The code for the PHASE project is available at: https://github.com/wuqinhua/PHASE and will be updated regularly during the submission process.

## Data availability

All data used in this study are available online from public studies. The COVID-19_Arunachalam_2020 data used in this study is available in the GEO database under accession code <u>GSE155673</u>^13^. The COVID-19_COMBAT_2022 dataset can be accessed through the EGA database with the code <u>EGAS00001005493</u>^14^. The COVID-19_Combes_2021 data can be accessed on the GEO database under accession code <u>GSE163668</u>^15^. The COVID-19_Liu_2021 data can be accessed on the GEO database under accession code <u>GSE161918</u>^16^. The COVID-19_Ren_2021 data can be accessed on the GEO database under accession code <u>GSE158055</u>^17^. The COVID-19_Schulte-Schrepping_2020 data is available in the EGA database under accession code <u>EGAS00001004571</u>^18^. The COVID-19_Schuurman_2021 data can be accessed on the GEO database under accession code <u>GSE164948</u>^19^. The COVID-19_Stephenson_2021 data is available in the EBI database under accession code <u>E-MTAB-</u> <u>10026</u>^20^. The COVID-19_Unterman_2022 data can be accessed on the GEO database under accession code <u>GSE155223</u>^21^. The COVID-19_Wilk_2021 data can be accessed on the GEO database under accession code <u>GSE174072</u>^22^. The COVID-19_Zhao_2021 data is available in the CNGB database under project code <u>CNP0001250</u>^23^. The COVID-19_Zhu_2020 data is available in the CNGB database under project code <u>CNP0001102</u>^24^. The Age_Terekhova_2023 data can be accessed on the Synapse database under accession code <u>syn49637038</u>^45^. The Age_Zhu_2023 data is available in the GEO database under accession code <u>GSE213516</u>^35^.

## Authors’ contributions

Y.L., J.L., and L.H. supervised the project. Q.W., J.L., and J.D. designed the models. Q.W., J.L., and J.D. developed the code of the model and performed benchmark analysis for model evaluation. J.L., Q.W., wrote the manuscripts. All other authors contributed to the design and implementation of this model, and read and approved the final manuscript.

## Acknowledgments

We thank technical support from the Data Science Platform of Guangzhou National Laboratory and the Bio-medical Big Data Operating System (Bio-OS).

## Declarations

Ethics approval and consent to participate

Not applicable

## References

1 Wen, L. et al. Single-cell technologies: From research to application. Innovation 3 (2022). 10.1016/j.xinn.2022.100342

2 Heumos, L. et al. Best practices for single-cell analysis across modalities. Nat Rev Genet 24, 550–572 (2023). 10.1038/s41576-023-00586-w

3 Chazarra-Gil, R., van Dongen, S., Kiselev, V. Y. & Hemberg, M. Flexible comparison of batch correction methods for single-cell RNA-seq using BatchBench. Nucleic Acids Res 49, e42 (2021). 10.1093/nar/gkab004

4 He, B. et al. CloudPred: Predicting Patient Phenotypes From Single-cell RNA-seq. *Pacific Symposium on Biocomputing*. Pacific Symposium on Biocomputing 27, 337–348 (2022).

5 Zeng, F., Kong, X., Yang, F., Chen, T. & Han, J. (2022). 10.1101/2022.06.20.496916

6 Mao, Y. et al. Phenotype prediction from single-cell RNA-seq data using attention-based neural networks. Bioinformatics 40 (2024). 10.1093/bioinformatics/btae067

7 Zhao, J. et al. Detection of differentially abundant cell subpopulations in scRNA-seq data. Proceedings of the National Academy of Sciences of the United States of America 118 (2021). 10.1073/pnas.2100293118

8 Dann, E. et al. Precise identification of cell states altered in disease using healthy single-cell references. Nat Genet 55, 1998–2008 (2023). 10.1038/s41588-023-01523-7

9 Ilse, M., Tomczak, J. & Welling, M. in International conference on machine learning. 2127-2136 (PMLR).

10 Rymarczyk, D., Borowa, A., Tabor, J. & Zielinski, B. in Proceedings of the IEEE/CVF Winter Conference on Applications of Computer Vision. 1721–1730.

11 Sundararajan, M., Taly, A. & Yan, Q. in International conference on machine learning. 3319–3328 (PMLR).

12 Dominguez Conde, C., et al. Cross-tissue immune cell analysis reveals tissue-specific features in humans. Science 376, eabl5197 (2022). 10.1126/science.abl5197

13 Prabhu S. Arunachalam1*, F. W., Chris Ka Pun Mok2*, Ranawaka A. P. M. Perera3*, Madeleine Scott1,4†, Thomas Hagan1†, Natalia Sigal1†, Yupeng Feng1†, Laurel Bristow5,, Owen Tak-Yin Tsang6, D. W., John Coller7, Kathryn L. Pellegrini8, Dmitri Kazmin1, Ghina Alaaeddine5, Wai Shing Leung6, Jacky Man Chun Chan6, Thomas Shiu Hong Chik6,, Chris Yau Chung Choi6, C. H., Michele Paine McCullough5, Huibin Lv2, Evan Anderson9, Srilatha Edupuganti5, Amit A. Upadhyay8, Steve E. Bosinger8,10, Holden Terry Maecker1, & Purvesh Khatri1, Nadine Rouphael5, Malik Peiris2,3, Bali Pulendran1,11,12‡. Systems biological assessment of immunity to mild versus severe COVID-19 infection in humans. Science (2020).

14, C. O.-M.-o. B. A. C. E. a. & Consortium, C. O.-M.-o. B. A. A blood atlas of COVID-19 defines hallmarks of disease severity and specificity. Cell 185, 916–938 e958 (2022). 10.1016/j.cell.2022.01.012

15 Combes, A. J. et al. Global absence and targeting of protective immune states in severe COVID-19. Nature 591, 124–130 (2021). 10.1038/s41586-021-03234-7

16 Liu, C. et al. Time-resolved systems immunology reveals a late juncture linked to fatal COVID-19. Cell 184, 1836–1857 e1822 (2021). 10.1016/j.cell.2021.02.018

17 Ren, X. W. et al. COVID-19 immune features revealed by a large-scale single-cell transcriptome atlas. Cell 184, 1895-+ (2021). 10.1016/j.cell.2021.01.053

18 Schulte-Schrepping, J. et al. Severe COVID-19 Is Marked by a Dysregulated Myeloid Cell Compartment. Cell 182, 1419–1440 e1423 (2020). 10.1016/j.cell.2020.08.001

19 Schuurman, A. R. et al. Integrated single-cell analysis unveils diverging immune features of COVID-19, influenza, and other community-acquired pneumonia. Elife 10 (2021). 10.7554/eLife.69661

20 Stephenson, E. et al. Single-cell multi-omics analysis of the immune response in COVID-19. Nat Med 27, 904–916 (2021). 10.1038/s41591-021-01329-2

21 Unterman, A. et al. Single-cell multi-omics reveals dyssynchrony of the innate and adaptive immune system in progressive COVID-19. Nat Commun 13, 440 (2022). 10.1038/s41467-021-27716-4

22 Wilk, A. J. et al. Multi-omic profiling reveals widespread dysregulation of innate immunity and hematopoiesis in COVID-19. J Exp Med 218 (2021). 10.1084/jem.20210582

23 Zhao, X. N. et al. Single-cell immune profiling reveals distinct immune response in asymptomatic COVID-19 patients. Signal Transduct Target Ther 6, 342 (2021). 10.1038/s41392-021-00753-7

24 Zhu, L. et al. Single-Cell Sequencing of Peripheral Mononuclear Cells Reveals Distinct Immune Response Landscapes of COVID-19 and Influenza Patients. Immunity 53, 685–696 e683 (2020). 10.1016/j.immuni.2020.07.009

25 He, B. et al. CloudPred: Predicting Patient Phenotypes From Single-cell RNA-seq. Pac Symp Biocomput 27, 337–348 (2022).

26 Zaheer, M. et al. Deep sets. Advances in neural information processing systems 30 (2017).

27 Silvin, A. et al. Elevated calprotectin and abnormal myeloid cell subsets discriminate severe from mild COVID-19. Cell 182, 1401–1418. e1418 (2020).

28 Markovic, S. S. et al. Galectin-1 as the new player in staging and prognosis of COVID-19. Scientific Reports 12, 1272 (2022).

29 Bojkova, D. et al. Targeting the pentose phosphate pathway for SARS-CoV-2 therapy. Metabolites 11, 699 (2021).

30 Stephenson, E. et al. Single-cell multi-omics analysis of the immune response in COVID-19. Nature medicine 27, 904–916 (2021).

31 Li, T. et al. Platelets mediate inflammatory monocyte activation by SARS-CoV-2 spike protein. The Journal of clinical investigation 132 (2022).

32 Merad, M. & Martin, J. C. Pathological inflammation in patients with COVID-19: a key role for monocytes and macrophages. Nature reviews immunology 20, 355–362 (2020).

33 Giamarellos-Bourboulis, E. J. et al. Complex immune dysregulation in COVID-19 patients with severe respiratory failure. Cell host & microbe 27, 992–1000. e1003 (2020).

34 Terekhova, M. et al. Single-cell atlas of healthy human blood unveils age-related loss of NKG2C+ GZMB− CD8+ memory T cells and accumulation of type 2 memory T cells. Immunity 56, 2836–2854. e2839 (2023).

35 Zhu, H. et al. Human PBMC scRNA-seq–based aging clocks reveal ribosome to inflammation balance as a single-cell aging hallmark and super longevity. Sci Adv 9, eabq7599 (2023).

36 Harries, L. W. et al. Human aging is characterized by focused changes in gene expression and deregulation of alternative splicing. Aging cell 10, 868–878 (2011).

37 Karagiannis, T. T. et al. Multi-modal profiling of peripheral blood cells across the human lifespan reveals distinct immune cell signatures of aging and longevity. EBioMedicine 90 (2023).

38 Nahata, M. et al. Bcl-2-dependent autophagy disruption during aging impairs amino acid utilization that is restored by hochuekkito. npj Aging and Mechanisms of Disease 7, 13 (2021).

39 Wagner, J. et al. A Single-Cell Atlas of the Tumor and Immune Ecosystem of Human Breast Cancer. Cell 177, 1330–1345 e1318 (2019). 10.1016/j.cell.2019.03.005

40 Miao, Y. et al. Adaptive Immune Resistance Emerges from Tumor-Initiating Stem Cells. Cell 177, 1172–1186 e1114 (2019). 10.1016/j.cell.2019.03.025

41 Litinetskaya, A. et al. Multimodal weakly supervised learning to identify disease-specific changes in single-cell atlases. bioRxiv, 2024.2007. 2029.605625 (2024).

42 Chen, C.-L. et al. An annotation-free whole-slide training approach to pathological classification of lung cancer types using deep learning. Nature communications 12, 1193 (2021).

43 Wolf, F. A., Angerer, P. & Theis, F. J. SCANPY: large-scale single-cell gene expression data analysis. Genome Biol 19, 15 (2018). 10.1186/s13059-017-1382-0

44 Xu, C. et al. Automatic cell-type harmonization and integration across Human Cell Atlas datasets. Cell 186, 5876–5891 e5820 (2023). 10.1016/j.cell.2023.11.026

45 Terekhova, M. et al. Single-cell atlas of healthy human blood unveils age-related loss of NKG2C(+)GZMB(-)CD8(+) memory T cells and accumulation of type 2 memory T cells. Immunity 56, 2836–2854 e2839 (2023). 10.1016/j.immuni.2023.10.013

