## Supplementary material for "Exploring phenotype-related single-cells through attention-enhanced representation learning": PHASE_Supplementary_Figure.docx

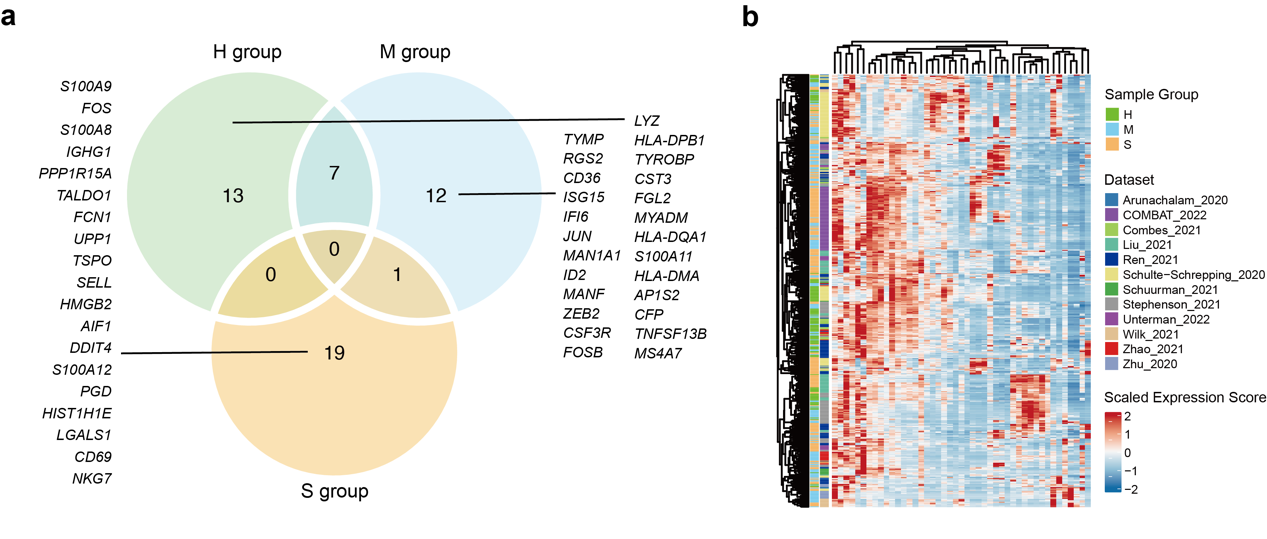
**Supplementary Fig. 1. Related exploration of the** **top-20 genes with gene attribution scores. a**, Venn plot of top-20 genes with gene attribution scores across three phenotype groups. **b**, Heatmap of the gene expression levels for the selected genes in (d) across individual samples, labeled by sample group and dataset on the left. The heatmap was clustered by hierarchical clustering.

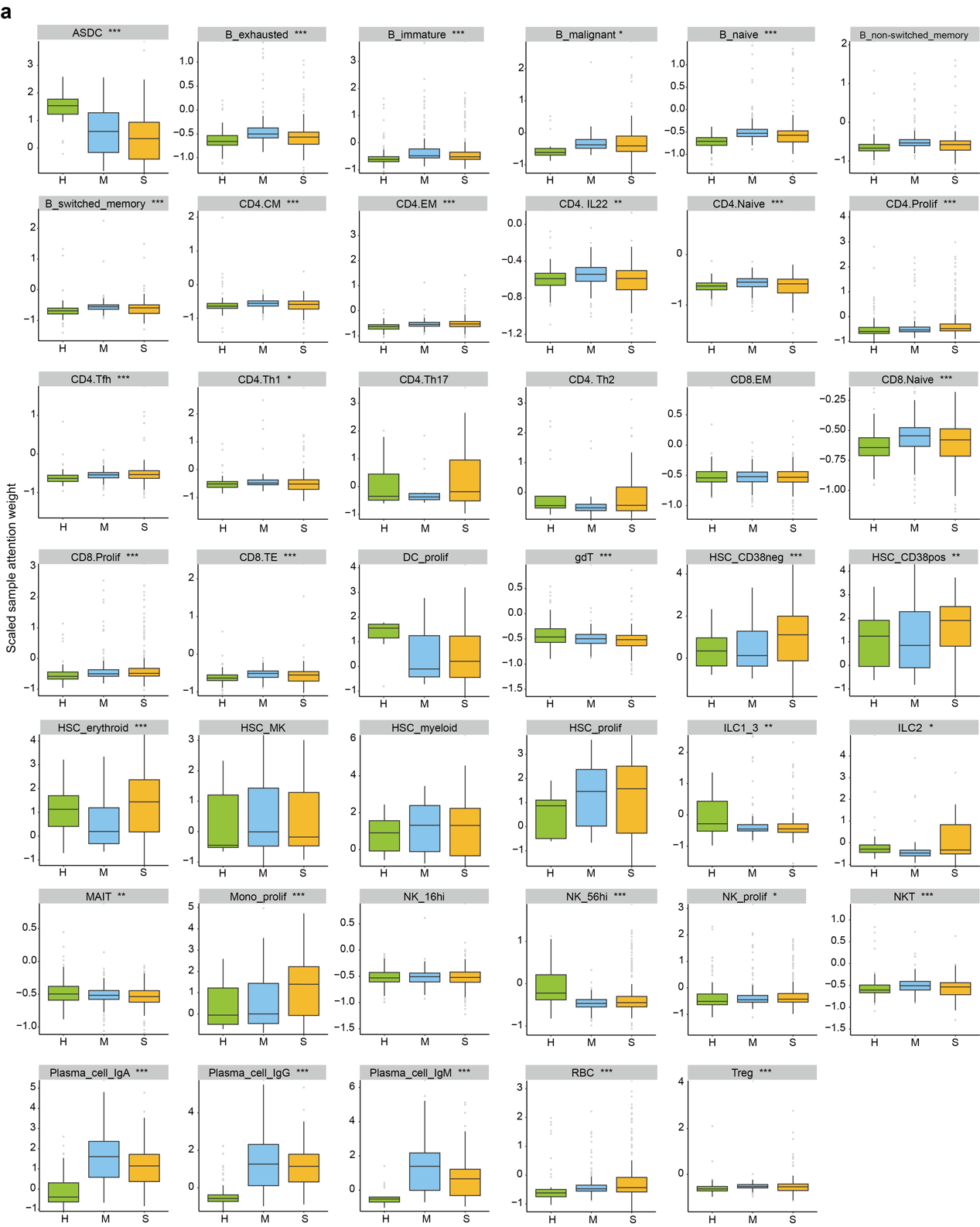
**Supplementary Fig. 2.** Comparison of scaled sample attention weights within individual cell subsets across sample groups. Boxplots represent the median, upper quartile, and lower quartile values for each group. The Kruskal-Wallis test was used to assess statistical significance. The statistical levels indicated as follows: *p ≤ 0.05, **p ≤ 0.01, ***p ≤ 0.001.

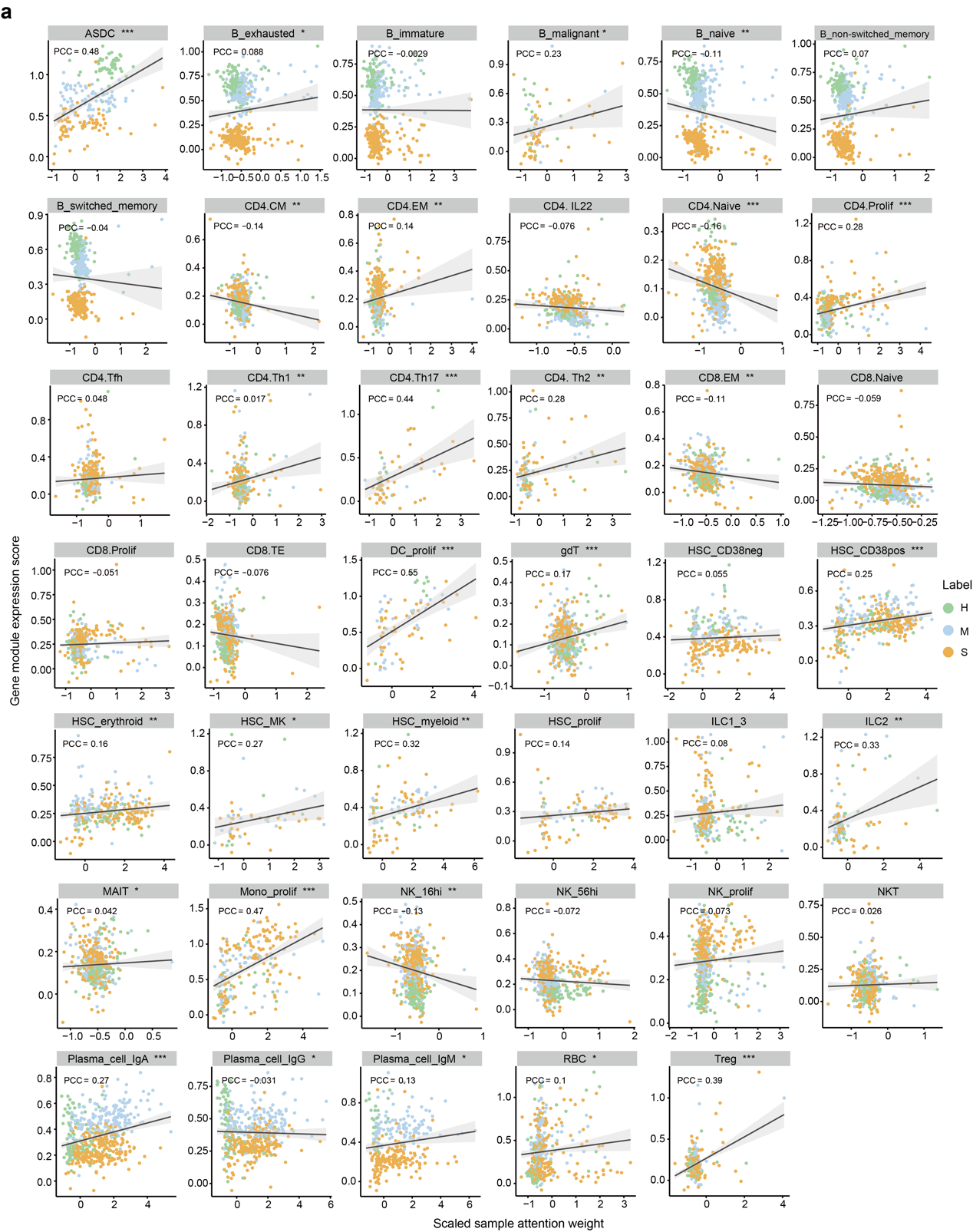
**Supplementary Fig. 3.** Pearson’s correlation analysis between the sample gene module expressions of group-specific high attributed genes and the scaled sample attention weights. The plot is labeled with correlation coefficient, a regression line and its confidence interval. The statistical levels indicated as follows: *p ≤ 0.05, **p ≤ 0.01, ***p ≤ 0.001.

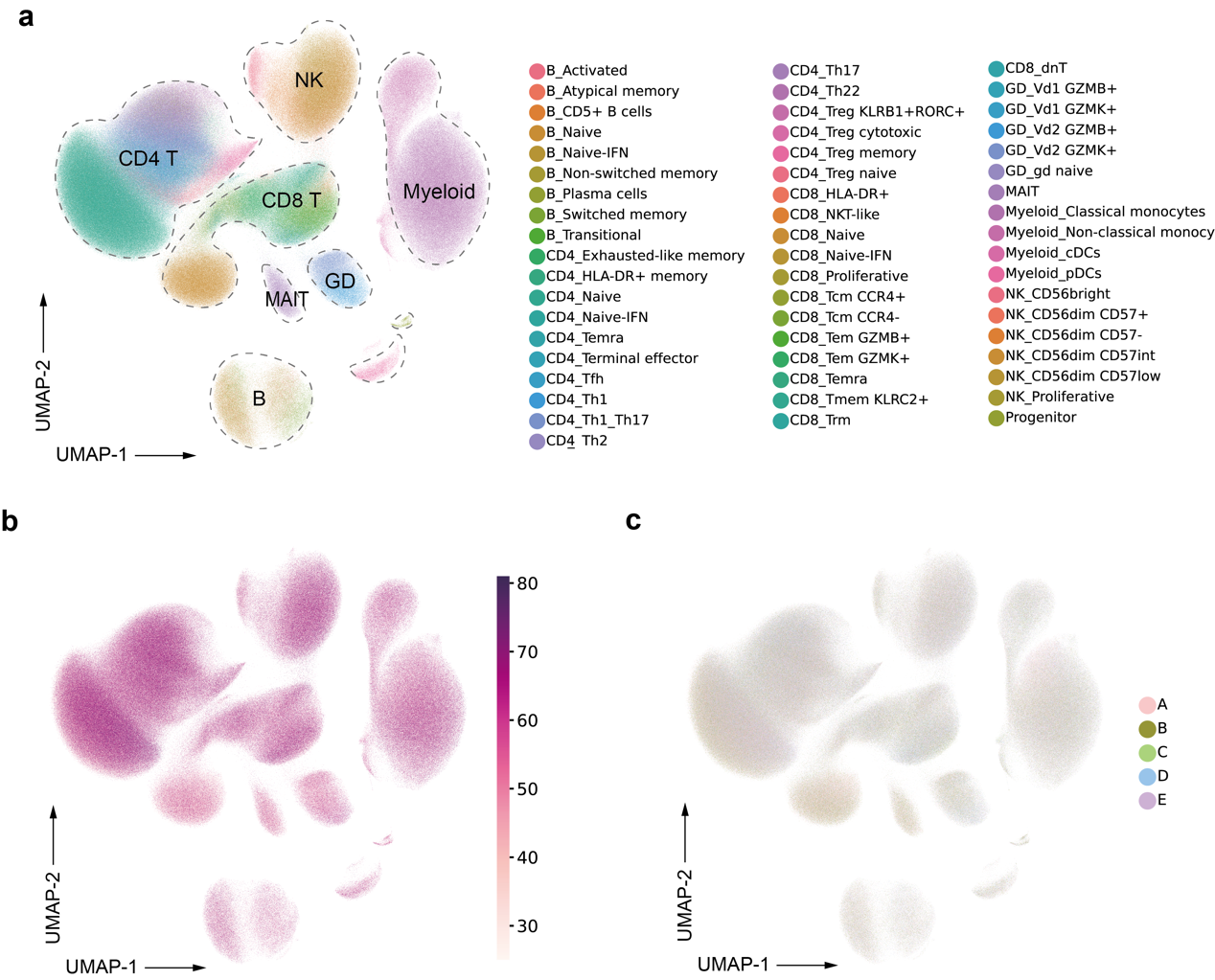
**Supplementary Fig. 4. UMAP of** **Age_Terekhova_2023 dataset. a**, UMAP plot of single cells from the Age_Terekhova_2023 dataset, colored by automatically annotated cell subsets, and labeled with cell subset ID and major cell subtypes. **b**, UMAP plot of single cells i from the Age_Terekhova_2023 dataset, colored by age. **c**, UMAP plot of single cells from the Age_Terekhova_2023 dataset, colored by age group.

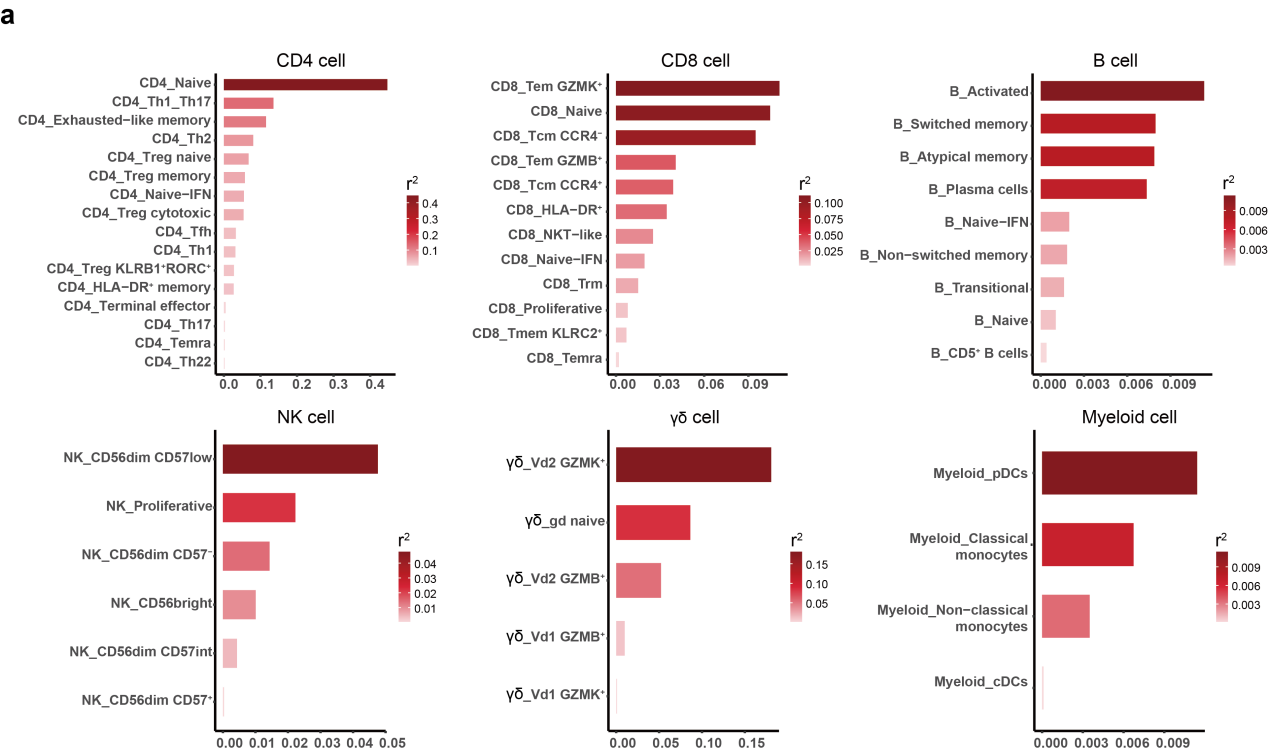
**Supplementary Fig. 5.** The R² values between z-score-scaled relative attention scores and age labels for each cell subset are computed and sorted by major cell type**.**

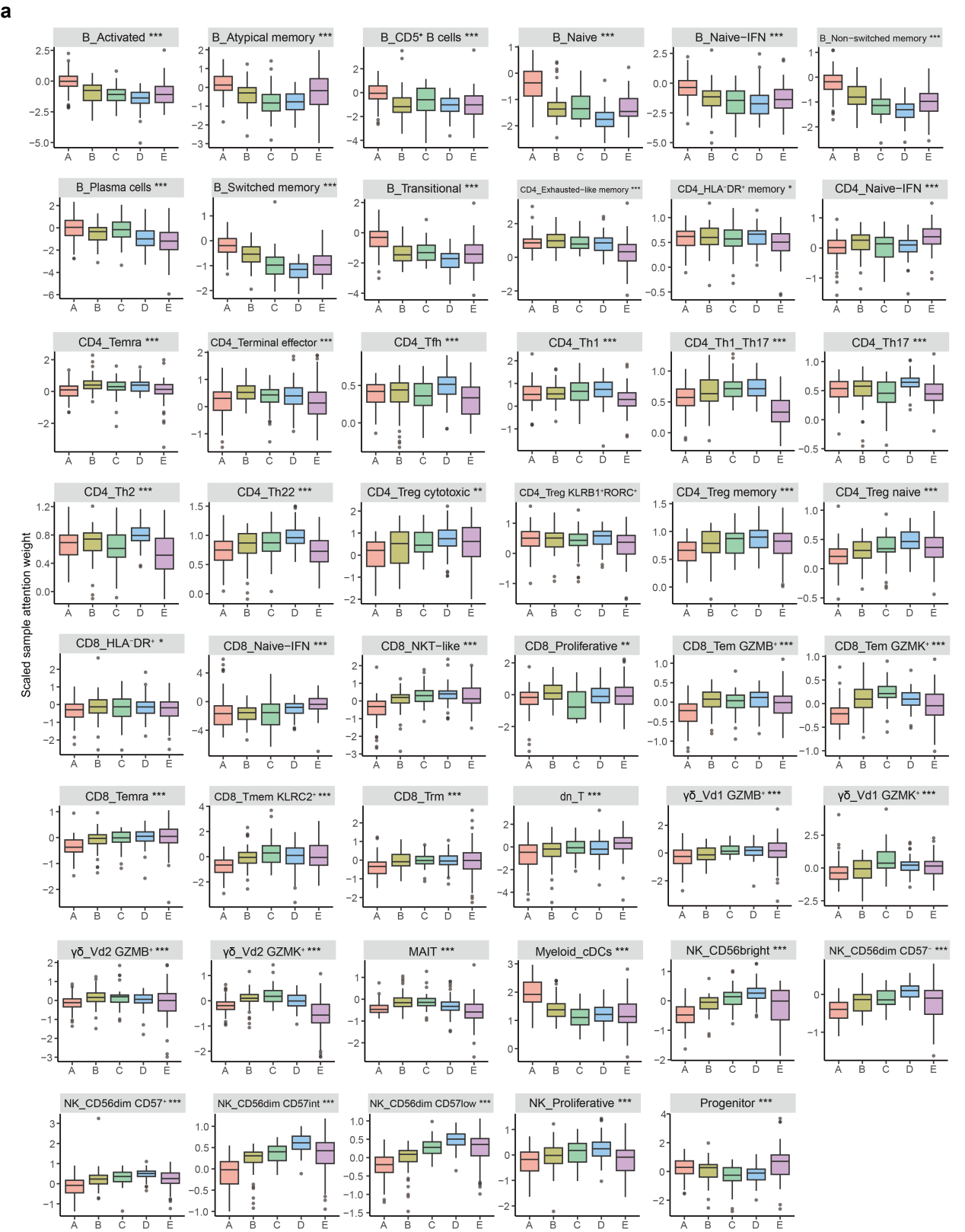
**Supplementary Fig. 6.** Comparison of scaled sample attention weights within individual cell subsets across age groups. Boxplots represent the median, upper quartile, and lower quartile values for each group. The Kruskal-Wallis test was used to assess statistical significance. The statistical levels indicated as follows: *p ≤ 0.05, **p ≤ 0.01, ***p ≤ 0.001.

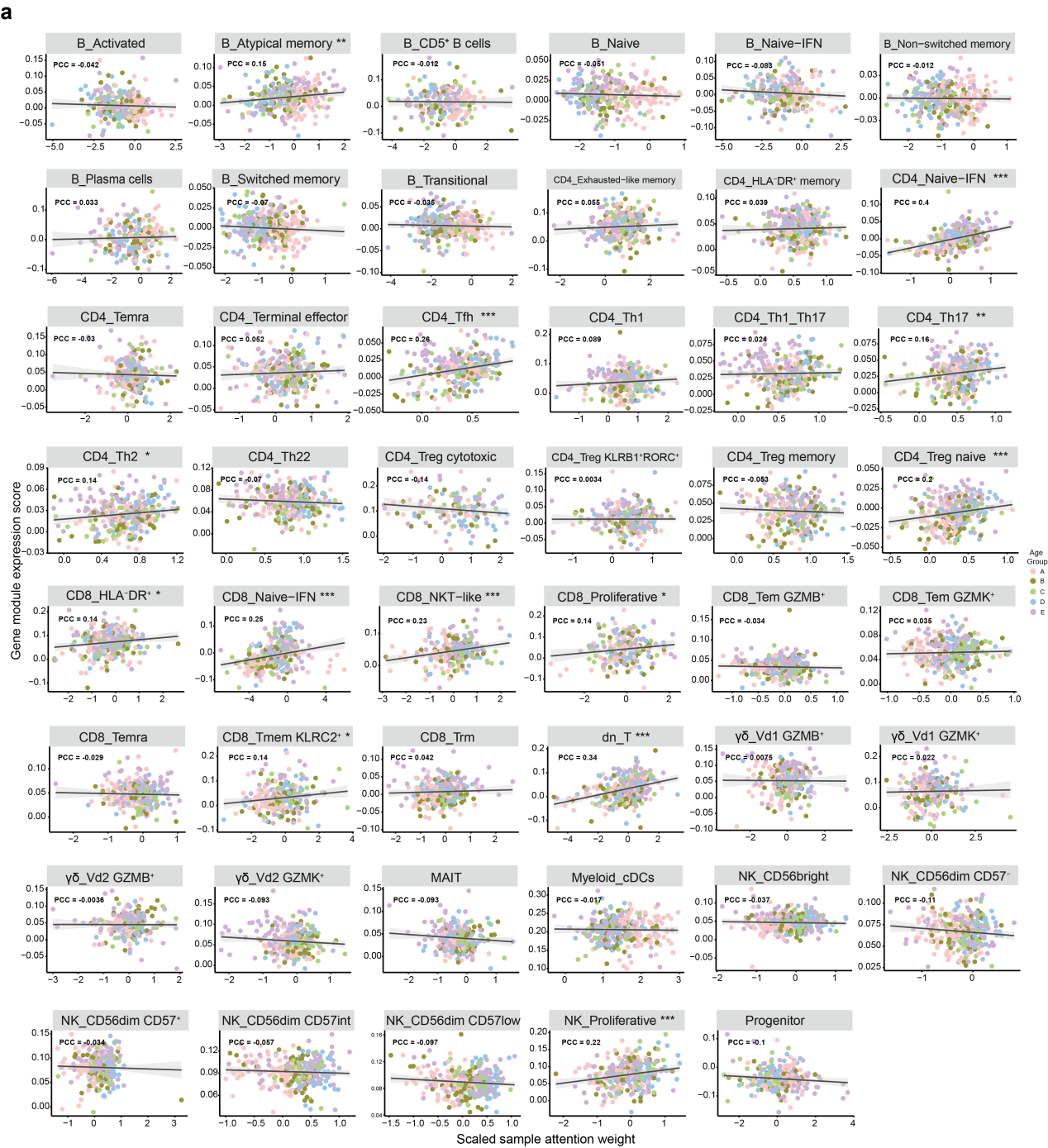
**Supplementary Fig. 7.** Pearson’s correlation analysis between the sample gene module expressions of group-specific high attributed genes and the scaled sample attention weights. The plot is labeled with correlation coefficient, a regression line and its confidence interval. The statistical levels indicated as follows: *p ≤ 0.05, **p ≤ 0.01, ***p ≤ 0.001.
